# Microbial Similarity Between Students in a Common Dormitory Environment Reveals the Forensic Potential of Individual Microbial Signatures

**DOI:** 10.1101/620948

**Authors:** Miles Richardson, Neil Gottel, Jack A Gilbert, Simon Lax

## Abstract

The microbiota of the built environment is an amalgamation of both human and environmental sources. While human sources have been examined within single-family households or in public environments, it is unclear what effect a large number of cohabitating people have on the microbial communities of their shared environment. We sampled the public and private spaces of a college dormitory, disentangling individual microbial signatures and their impact on the microbiota of common spaces. We compared multiple methods for marker gene sequence clustering, and found that Minimum Entropy Decomposition (MED) was best able to distinguish between the microbial signatures of different individuals, and was able to uncover more discriminative taxa across all taxonomic groups. Further, weighted UniFrac- and random forest-based graph analyses uncovered two distinct spheres of hand or shoe associated samples. For hand-associated samples, connection between cliques was enriched for hands, implicating them as a primary means of transmission. By contrast, shoe-associated samples were found to be freely interacting, with individual shoes more connected to each other than to the floors they interact with. Individual interactions were highly dynamic, with groups of samples originating from individuals clustering freely with other individuals, while all floor and shoe samples consistently clustered together.

**Importance:** Humans leave behind a microbial trail, regardless of intention. This may allow for the identification of individuals based on the ‘microbial signatures’ they shed in built environments. In a shared living environment, these trails intersect, and through interaction with common surfaces may become homogenized, potentially confounding our ability to link individuals to their associated microbiota. We sought to understand the factors that influence the mixing of individual signatures, and how best to process sequencing data to best tease apart these signatures.

## Introduction

Numerous recent studies have uncovered the extent to which humans influence the microbial ecology of the spaces they occupy through microbial exchange between skin and the built environment. Most of these studies have focused on home-associated microbial communities(1–3), with home size, number of occupants, and building materials differentiated between sampling locations. Each of those confounding factors may have significant impacts on microbial community structure, and they are difficult to disentangle. Other studies have focused instead on the microbial ecology of public spaces, such as classrooms and hospital entrance halls(4–8). Although they have been able demonstrate that most of the taxa colonizing those spaces are skin-associated, they are unable to link individual human microbial signatures to their data.

Microbial flow in the built environment is a keen topic of interest. Cohabitation of multiple individuals has been shown to influence the microbiota of common spaces, and of the constituents themselves(1, 7, 9). Common areas may also serve as sites of exchange between individuals, with implications for disease control. Also unclear are how methodological differences in sequence clustering impact the ability of these studies to link individuals to their surroundings through microbial similarity.

Dorm buildings, which have a standardized architectural design, common building materials and furnishings between rooms, and even a common ventilation system, represent an intriguing model system in which to characterize the direct effects of an individual’s skin microbiota on their surroundings, and to further elucidate the forensic potential of skin microbial signatures. Individuals shed around 30 million bacterial cells per hour(10), and thus leave behind a “microbial fingerprint”. In one sense, dorm rooms represent a number of identical replicates that can be used to uncover general patterns of human microbial exchange with the built environment. In a different sense, they are a “metacommunity” in which it is possible to record a network of interaction by logging visits between rooms and the use of common spaces. The divide between private rooms and common spaces such as hallways, lounges, and restrooms further enables us to tease apart individual microbial signatures in shared spaces.

To explore the divide between public and private, we sampled 37 participants from the University of Chicago’s eight floor South Campus residence hall, with four timepoints over 3 months. Participants were drawn from one “house” in the dormitory, which serves a subset of the dormitory floor plan with shared common space and bathrooms. From participants, we swabbed both skin sites, such as hands, and personal effects, such as bed sheets. Additionally, common surfaces such as tables and bathrooms were also sampled. Together, this collection of surfaces encompasses the divide between private and public space in the dormitory.

To determine how to optimize the inference of individual microbial signatures, we employed three sequence processing methods to find the most discriminative in characterizing individuals. It has been observed that in many built environment studies, a large fraction of reads were attributed to a small number of OTUs(1, 9). These OTUs come from a small selection of skin-associated taxonomic groups, including corynebacteria, staphylococci, pseudomonads, and streptococci. As much of the differentiation between individuals occurs within a small number of taxonomic groups, it is unclear how to optimize sequence clustering for forensic inference as OTU clustering may lump together similar sequences by design. UPARSE(11) is an greedy OTU clustering algorithm for sequence processing, relying on a greedy clustering method that uses highly abundant sequences as seeds for clustering. Sequence based methods do not employ OTU clustering and provide a higher resolution for sequence differentiation. DADA2(12) is a reference-free sequence based algorithm that partitions reads based on an error model generated from the dataset. Minimum Entropy Decomposition(13) (MED) is an unsupervised version of oligotyping(14), a method that derives sequences by looking for regions of high Shannon entropy among 16S regions, and decomposing them into constituent oligotypes. Oligotyping has been used to explore variation in host associated bacteria, such as in *Blautia* found in sewage systems(15).

## Results

### Clustering Methodology Impacts the Success of Forensic Inference

Each of the sequence processing methods produced a different picture of the microbial diversity of the dormitory. UPARSE recovered the largest number of distinct sequences (6011) along with the greatest number of phyla. MED recovered fewer sequences (3353), and fewer phyla (9), but recovered more members within each phylum (**Supplementary Table 1**). DADA2 recovered nearly the same phylum level diversity as UPARSE (23 vs 25), but fewer sequences (4307). MED also had a significantly smaller phylogenetic distance between taxa (Wilcoxan Rank-sum test, p < 2.2e-16) than both DADA2 and UPARSE (**Figure 1a),** indicating that MED recovered much more closely related sequences.

**Figure 1:**
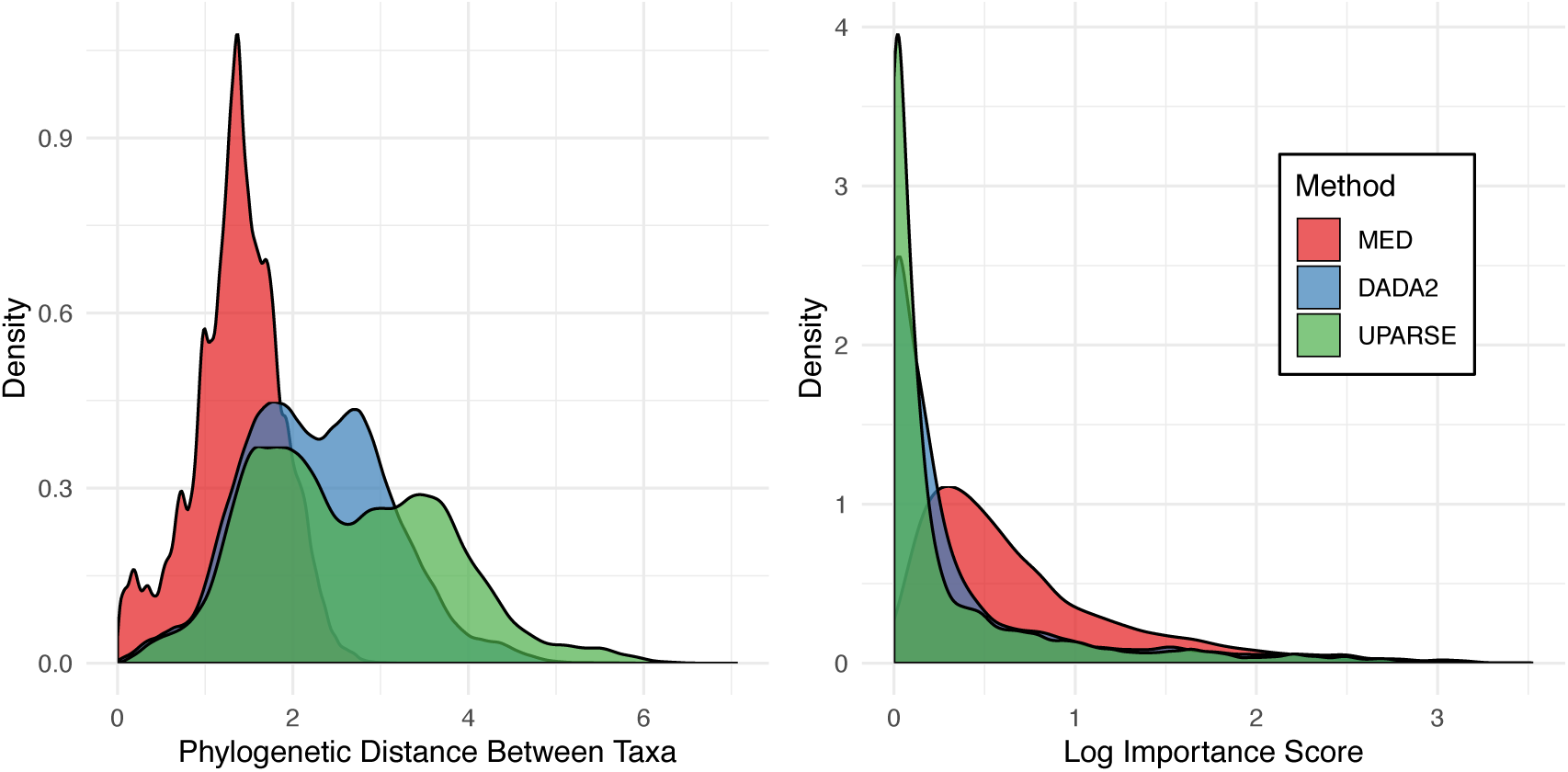
(a) The distribution of phylogenetic distance between all taxa in each sequence processing method. MED recovers more highly related taxa than DADA2 or UPARSE. (b) The distribution of importance scores over all taxa, grouped by sequence processing method. The y-axis is log-transformed to aid visualization.

Since we were most interested in classifying individuals, we compared each method using a random forest trained on surfaces that closely associate with the hands of only one individual, in order to test their forensic inference. There is a major divide between floor and hand-associated samples (**Figure 2).** Floor associated samples, including shoes and floors, inhabit a different space compared to hand associated samples, and this division significantly structures these communities (ANOSIM on Bray-Curtis Distance R=0.2821, P=0.001). Thus, to predict which individual’s hands a surface had interacted with, bed sheets, desks, and door handles of the participant rooms are most useful.

**Figure 2:**
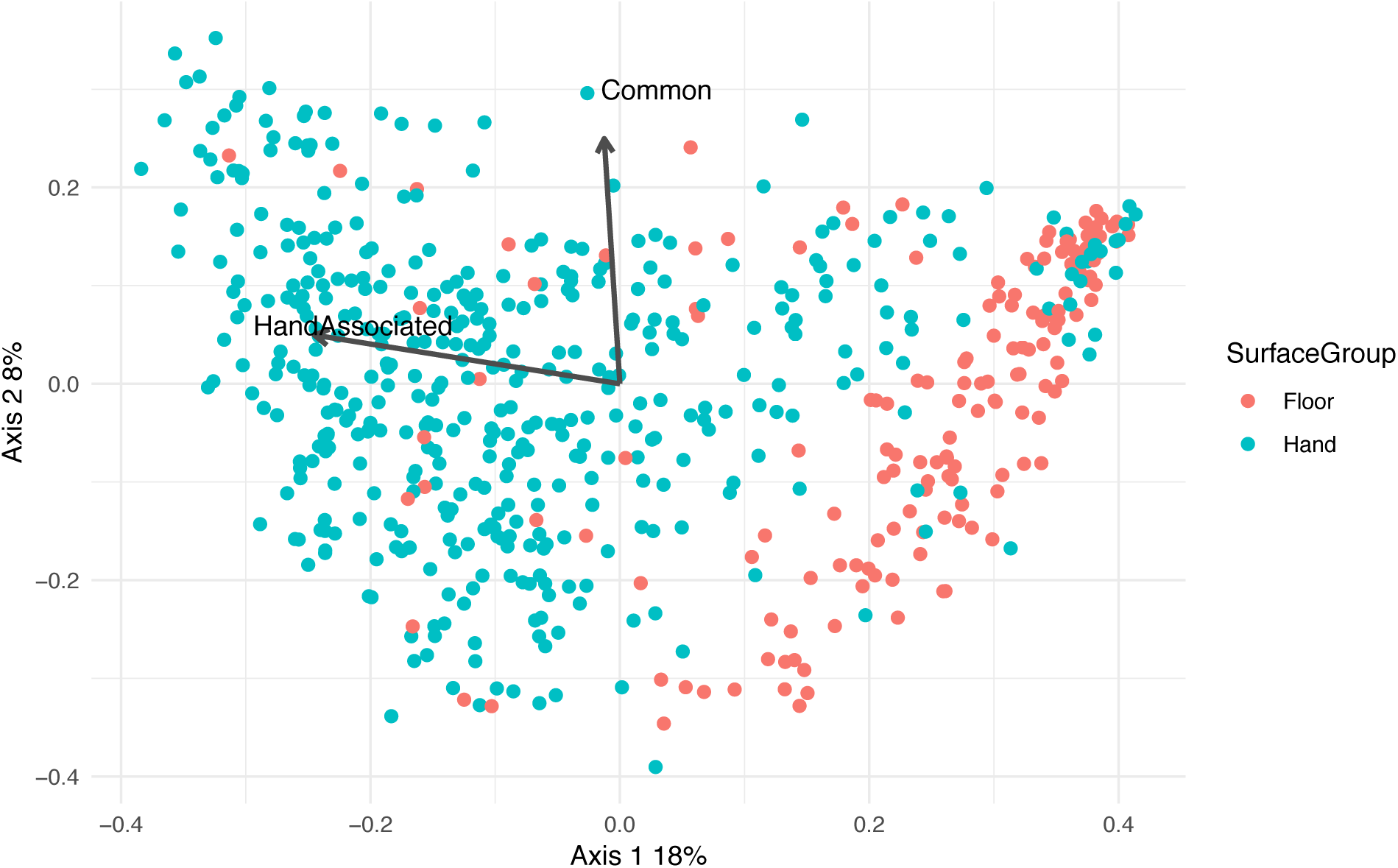
A principal components (PCoA) plot based upon the Bray-Curtis distance. Significant environmental vectors are plotted over the data.

These models were implemented using an random forest(39), which allows for the interrogation of similarity between samples. The model was then tested on hand samples from the same individuals, with the resulting accuracy summarized in **Table 1**. The standardized method of interpreting the success of classifiers is the error ration, which quantifies how well the random forest does at predicting the correct individual relative to the success expected by chance(40). An error ratio above two is commonly used as a significance threshold, and a higher ratio indicates better performance. All methods performed significantly better than random, but MED clearly outperformed UPARSE and DADA2 in our dataset. **Supplementary Figure 2** presents the confusion matrix generated by MED. Samples that fall on the diagonal are correctly classified by the random forest model. Most (79.57%) fall on the diagonal of the plot. However, for certain individuals, their hand samples are misclassified in every instance.

**Table 1:** Random Forest Accuracy and Error-Ratios

| Method | UPARSE | DADA2 | MED |
| --- | --- | --- | --- |
| Accuracy (CV-5) | 60.96% | 71.06% | 79.57% |
| Error-Ratio | 2.49 | 3.36 | 4.76 |

Interestingly, the largest source of classification error was the presence of roommates in the room. In fact, the classification error of an individual was linearly related to the number of roommates that individual had (R-squared 0.3143, P < 0.0001), with classification error increasing by 18 percentage points for each roommate. The relationship is shown in **Supplementary Figure 3**. The random forest model attempts to use differences in taxa abundance between individuals to classify individuals. If two individuals interact and exchange bacteria, differences in abundance decrease, which in turn increases model error. Roommates had a significantly smaller weighted UniFrac distance between them than individuals residing in different rooms. (Wilcoxon Rank-sum test, W = 409660000, p < 2.2 × 10^−16^)

### Classification of Individuals is Driven by Specific Taxa

The random forest model is able to rank individual sequences or OTUs by their importance in successful classification. As seen in **Figure 1**, there are differences in the distribution and average importance score across phyla, and all of these are significant to .05 by Kruskal-Wallis test. Furthermore, MED has a significantly higher importance score in all phyla that overlap between all three methods except for Cyanobacteria, Fusobacteria, and Deinococcus-Thermus (Wilcoxan Rank-Sum Test, FDR p < .05) (**Supplementary Figure 4**).

It has been noted that there are taxa indicative of different sexes.(41) To see if there were enriched taxa between men and women from room samples, we looked for differentially enriched taxa using DESeq2. The most significantly enriched taxon is *Lactobacillus iners*, an inhabitant of the female reproductive tract. Certain corynebacteria were also noted to be more abundant in men, as seen in **Supplementary Figure 5**. Using these enriched taxa, we used the random forest to predict whether a subject is a man or woman, with an error ratio of about 2.5.

### Metacommunity Structure

In addition to classifying individuals, we sought to recapitulate the geographical structure of the dorm using graphical models. To do this, we constructed a threshold graph of the weighted UniFrac distance between samples, with a threshold of 0.1. As seen in **Figure 3**, the dorm has two large subgraphs, along with a number of orphaned graphs. These two groups consist of floor-associated (shoes and floors) and hand-associated (hand, doorknob, bed, and desk) samples. The orphaned graphs are mostly samples from one individual. As expected, common surfaces in the hand-associated realm serve as an anchor for their subgraph, connecting a number of different people, while hallway floors serve the same role for individual shoes. By contrast, orphaned graphs appear to indicate the stability of an individual’s microbial signature over time and a lack of interaction with other samples.

**Figure 3:**
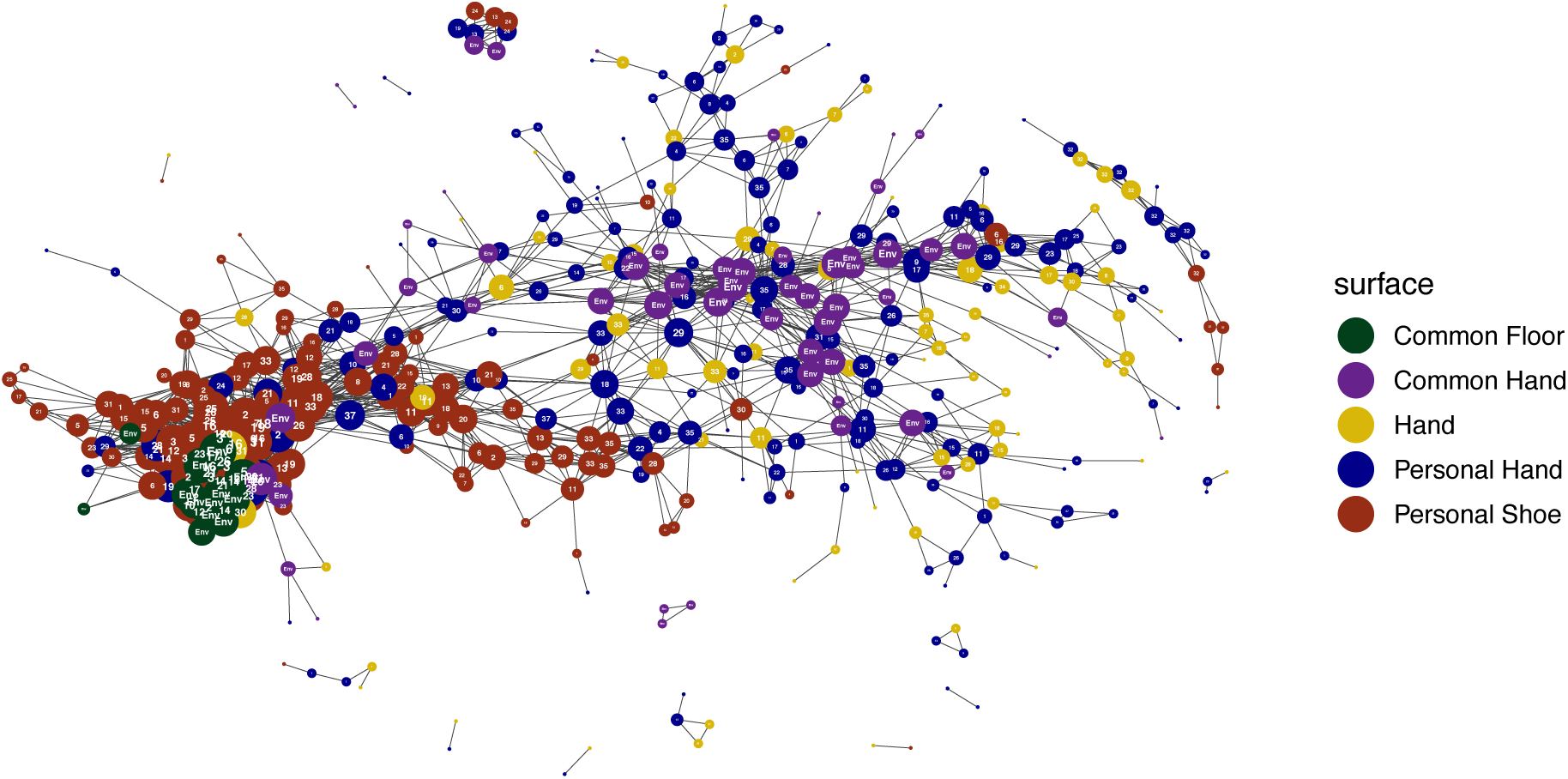
A Weighted-Unifrac graph of all samples, thresholded to be below .35 distance between individuals. Samples are colored by individual or environmental ID, while they are shaped by one of the four large sample types, personal vs common and if they are hand or show associated. Common hand-associated surfaces act as a scaffold, connecting between themselves, along with connecting many distinct individuals.

Further, we can also calculate the assortativity of different metadata criteria. Assortativity is a metric used to quantify how often a node attaches to a similar node, with higher assortativity reflecting higher connectivity between similar nodes. As seen in Table 2, sex and floor have the highest assortativity, while timepoint is the least important. This indicates that floor and sex are more important in generating the graph structure, and implies that the microbial signature of individual surfaces across the sampling period is stable.

**Table 2:**
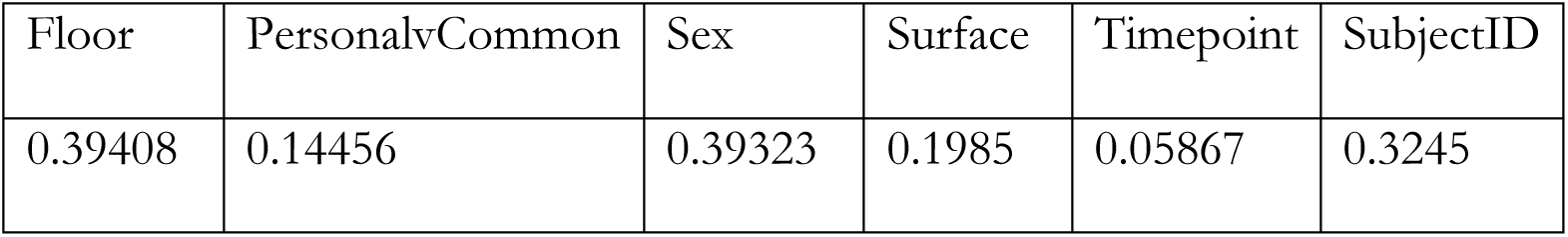
Assortativity of Metadata Factors

While a graph can be constructed using a beta-diversity metric (in our case weighted UniFrac distance) as above, the distance metric may not be sensitive to the microbial community of an individual. Since there is information to be gained from aggregating samples into a larger individual signature, we also constructed a graph using random forest proximity. The proximity values from the random forest are akin to a distance, and take into account the same signature used to classify individuals. It is also much sparser, as the random forest is trying to minimize distances between samples from the same individual, while keeping samples between individuals distinct. The resulting graph can be seen in **Supplementary Figure 6a and 6b**, where samples are colored by individual and surface type, respectively.

Graph based clustering analysis methods are often used in describing interactions in social networks. Using the Infomap clustering algorithm(42), which uses flow within a network to generate groupings, we looked at how bacterial exchange grouped our samples. The Infomap algorithm is also hierarchical(43), allowing for samples to inhabit “Top Modules” which are large scale groupings, and then submodules that indicate community clustering within top modules. Using this algorithm, we identified 8 top modules (**Figure 4)**, with module 1 encompassing almost all shoe and floor samples. Of particular interest were how samples grouped over time. Further, among surfaces connecting top modules, hands were significantly enriched, and other hand-associated surfaces showed enrichment, including common tables, doors, and bathroom doors. (**Supplementary Figure 7**) Since the dorm represents a multilayer graph, where each timepoint forms a distinct layer of interaction, we employed a multilayer implementation of the algorithm(44) to look at the stability of interactions over time. This is presented in **Figure 5**, where samples are clustered at each timepoint and their membership in clusters in tracked over time. Shoe and floor samples showed high stability over time, where most samples co-cluster over time in the same clusters (**Figure 5a**). Similarly, common surfaces had the same pattern, wherein common floor samples clustered consistently, while common hand samples could be affiliated with different samples (**Figure 5b)**. Individual samples, for example individual 1 and 29 (**Figure 5c**), showed the ability to cluster freely with other samples.

**Figure 4:**
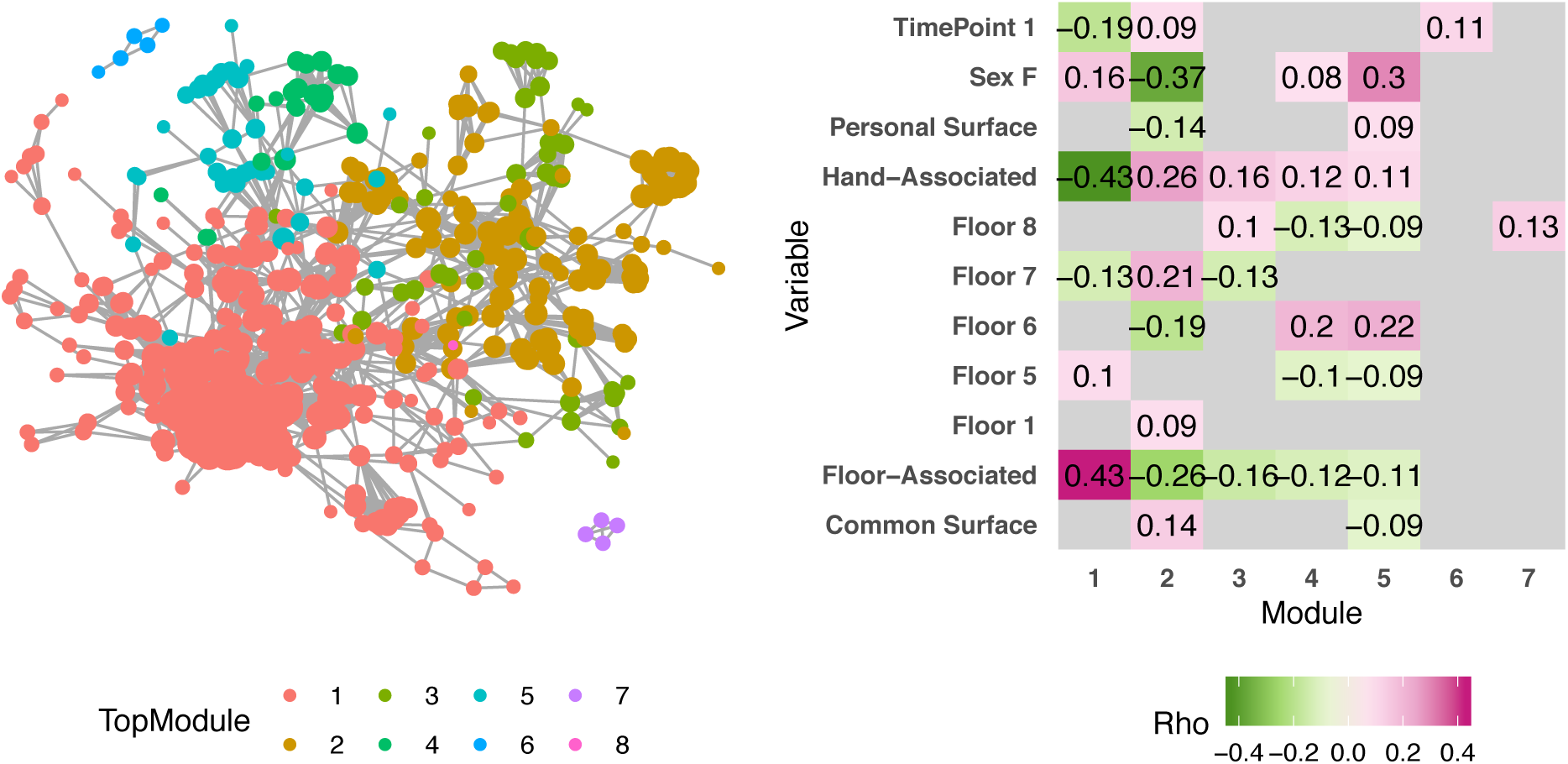
**(a)** A graph generated using random forest proximity scores, trained to distinguish individuals. It is thresholded by proximity greater than 0.076. It is colored by clique. Module 1 is mostly composed of shoe and floor samples, similarly to Figure 3. **(b)** Significant Spearman correlations (p<0.05) between each module and various metadata categories.

**Figure 5:**
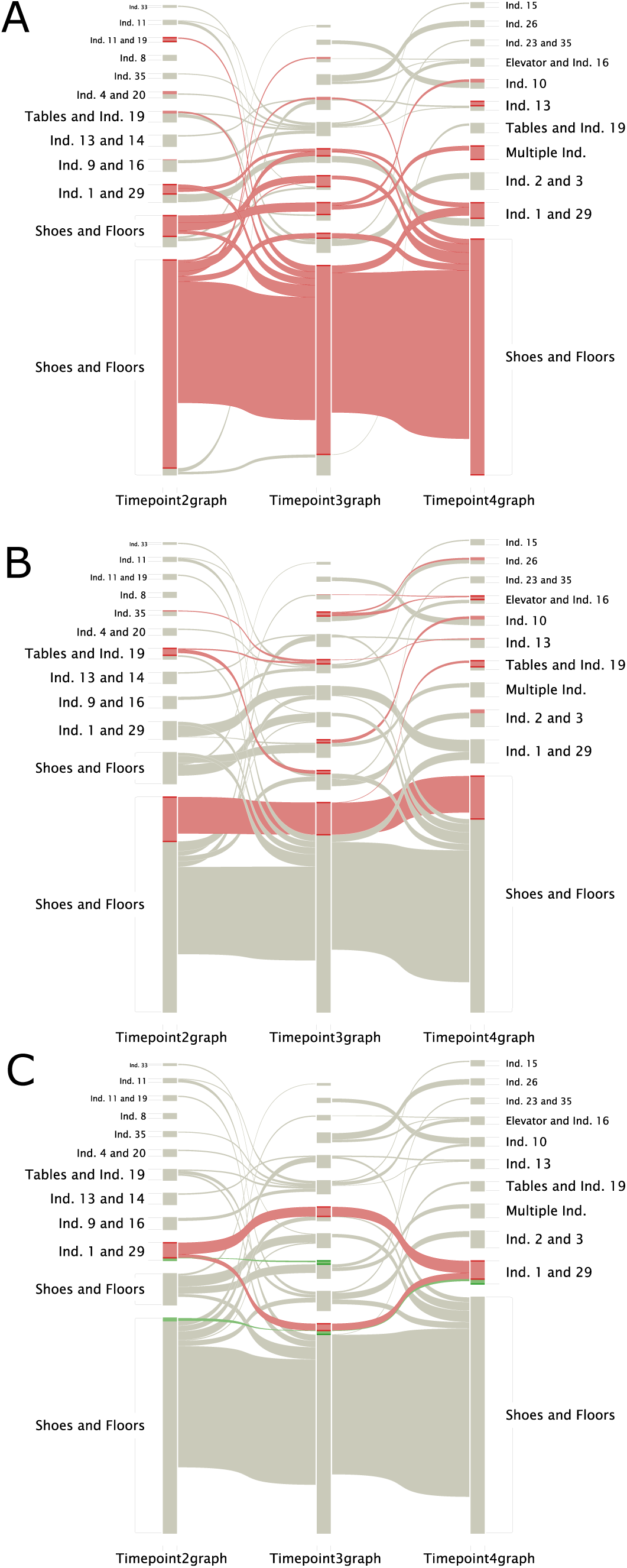
Alluvial Diagrams depicting the clustering of samples over time. (a) All samples that were floor-associated (hallway floors, bedroom floors, and shoes) were colored red. (b) By common surface. (c) Individual 1 (red) Individual 29 (green).

## Discussion

The use of human microbial signatures as trace evidence remains a young and inexact science. In order for this developing field to become a useful forensic tool, methods will need to be optimized and the myriad factors which influence our microbial interaction with built environments will need to be disentangled. Here, we compared classification methods to link residents to their rooms and personal effects in a common dormitory environment. For classifying individuals, Minimum Entropy Decomposition seems to be the best choice, but it appears that exact sequence variants in general are better at identifying individuals than OTU-clustering methods. This is unsurprising, as exact sequence variants maximize our insight into the microbial strains that differ between individuals that can be obscured by higher-level OTU-clustering. At the same time, this contrasts with observations that MED can produce incorrect sequence variants from mock communities. The advantage of MED seems to be that it is able to recover more diversity within the main skin-associated taxa from the phyla Proteobacteria, Fusobacteria, Bacteroidetes, Fusobacteria, and Actinobacteria. It is also able to recover higher importance scores even at the genus level, indicating that is it able to produce more individual-specific sequences within common skin taxa, as the importance score only measures the usefulness in classification between individuals.

The high accuracy of classification shows that skin associated samples, in particular bed sheets and door handles are useful in predicting the individuals inhabiting those rooms. Furthermore, samples within 2-4 weeks also appear useful in prediction, indicating that the signature is stable over long time scales. It is confounded by roommates, which is unsurprising, due to microbial exchange within one room.

This study explores the metacommunity structure of the college dormitory, which clusters into two distinct spheres-hand or shoe associated samples. Within each type, the arrangement between common and personal samples differs, with personal shoe associated samples freely associating, while hand associated sample often only associate between individuals when connected by a common surface. When examining graphs generated by random forest, the shoe associated structure remains the same, while individual signatures closely cluster with themselves. A multilayer graph reveals that individual signatures freely intermingle at different timepoints, while shoe and floor samples have large continuous interaction. This is likely a result of the high exchange between shoes and floors, which homogenizes the signature of shoe and floor samples.

## Materials and Methods

### Study Design and Sample Collection

We collected personal samples from 37 participants in 28 distinct dorm rooms (**Supplementary Table**). Samples were collected by swabbing a sterile cotton BD-Swube applicator against the dry surface of interest. Sampling kits were given to study participants for self-sampling with instructions. The desk, floor, fitted bed sheet, and interior doorknobs of each participant’s room, along with the dominant hand and shoe of the participant, were sampled at four timepoints. The first timepoint occurred before occupants left for a scheduled school break (end of a quarter) and then immediately upon return. The third and fourth timepoints were taken 2 and 4 weeks after spring break.

Participants also completed a questionnaire which collected basic information on the subject, the conditions specific to their dorm room, and who they interacted with in their dorm room during the sampling period. This questionnaire was completed each time a set of samples was collected.

Common surfaces were also sampled similarly. Common surfaces specific to the 5^th^ floor included tables in the dormitory lounge, and the handle of the entry door to the lounge. On each floor of the dormitory, the door handles of bathrooms, the floors of each hallway, and the elevator buttons were sampled. Each floor had its own unique combination, and these were swabbed at the same time as personal surfaces.

### Sample Processing

DNA was extracted from each sample using a low biomass variation of the MO BIO Powersoil DNA extraction protocol. 16s rRNA was amplified with the Earth Microbiome 16S Illumina Amplicon Protocol(16). The V4 region of the 16s rRNA gene was targeted with the 515F-806RB primer pair. Sequencing was performed using a Illumina Miseq sequencer with the protocol described in Caporaso et al. 2012(17).

### Sequence Processing

Each method was processed using the default workflows provided in reference papers.

## UPARSE

Demultiplexed sequences were merged using vsearch(18) with 10,040,708 successful paired end reads merged together. Sequences were quality filtered with a maximum expected error of 0.5, with 9,057,613 remaining sequences. Sequences were then dereplicated for 1,276,202 unique sequences. Sequences were then clustered at 97% identity, with 11658 OTUs and 42539 chimeras. Sequences were then matched to OTUs with 93.28% of sequences matched to OTUs. 6011 OTUs passed sequence processing. Chloroplast and mitochondrial DNA was removed, and samples were rarefied to 4000 counts per sample.

## MED

Sequences were processed according to the methods described in Meren et al 2015(13). Demultiplexed paired-end reads were merged using illumina-utils(19), with Q30 check imposed on sequences, leading to 10,023,266 successfully merged out of 10,023,266 reads. Gaps between sequences were padded with blanks, and samples were decomposed using a -M of 100. 1,732,615 outliers were removed by quality control, and remaining sequences were sorted into 3,748 nodes after refinement. 3,352 passed quality control. Chloroplast and mitochondrial DNA was removed, and samples were rarefied to 4000 counts per sample.

## DADA2

The filtering step of DADA2 was run with no ambiguous base (maxN of 0), maximum expected errors of 2, quality of truncation of 2. All other commands were run on default settings. Sequences were merged after performing quality filtering. After merging, 34043 sequences were observed, and 18329 sequences were not chimeras. 4307 unique sequences passed final quality filtering. Chloroplast and mitochondrial DNA was removed, and samples were rarefied to 4000 counts per sample.

### Taxonomic identification

All sequences were taxonomically identified using the same implementation of RDP(20) implemented in DADA2 to enable comparison between the sequencing methods. Taxonomy was assigned using the SILVA(21) training set version 123.

### Phylogenetic Trees

Sequences were aligned with the R package *MSA*(22), using the Muscle(23, 24) algorithm. Phylogenetic trees were then generated using the R package *Phangorn*(25). The tree was created by neighbor joining, and fitted with GTR model.

### Data Analysis and Visualization

Data cleaning and shaping was performed using R 3.3.2-R3.3.5 and the packages dplyr(26) reshape2(27). Visualization and analysis were performed using *phyloseq*(28), igraph(29), ggnetwork(30), and *ggplot2*(31). Random forests were generated using randomForest(32) and ranger(33). Differential abundance calculation were performed using DESeq2(34). Diversity measures were calculated using *vegan*(35). Inspiration was taken from Callahan et al. 2017(36). Community clustering was _performed_ using the InfoMap(37, 38) and alluvial diagrams generated using the “Map & Alluvial Generator” (http://www.mapequation.org/apps/MapGenerator.html).

## Acknowledgements

This work was sponsored by National Institutes of Justice award 2015-DN-BX-K430. Sophia Weaver for indispensable assistance in recruiting individuals. Study IRB Number: IRB15-0373. Approved by BSD IRB Committee A, The University of Chicago Biological Sciences Division/University of Chicago Medical Center.

---

**Supplementary Table 1:**
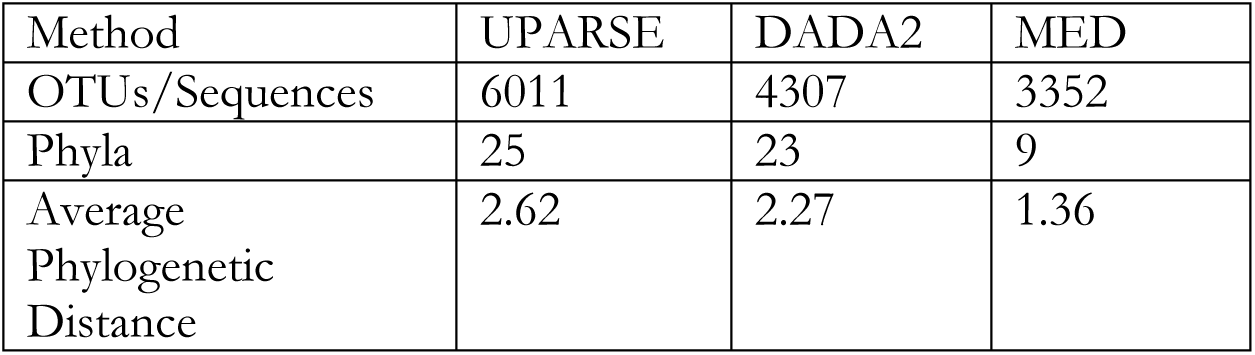
The OTU abundance and phylum-level diversity of each method. (fill this out with every taxonomic level.)

| Method | UPARSE | DADA2 | MED |
| --- | --- | --- | --- |
| OTUs/Sequences | 6011 | 4307 | 3352 |
| Phyla | 25 | 23 | 9 |
| Average Phylogenetic Distance | 2.62 | 2.27 | 1.36 |

**Supplementary Table 2:** Summary of the number of participants on each floor of the dormitory and which common surfaces were sampled on each floor.

| Floor | Number of Participants | Common Surfaces |
| --- | --- | --- |
| 5 | 11 | Male Bathroom Door Handle, Hallway Floor, Entry Door, Common Table, Elevator Buttons |
| 6 | 5 | Female Bathroom Door Handle, Hallway Floor, Elevator Button |
| 7 | 9 | Male Bathroom Door Handle, Hallway Floor, Elevator Button |
| 8 | 12 | Female Bathroom Door Handle, Hallway Floor, Elevator Button |

**Supplementary Figure 1:**
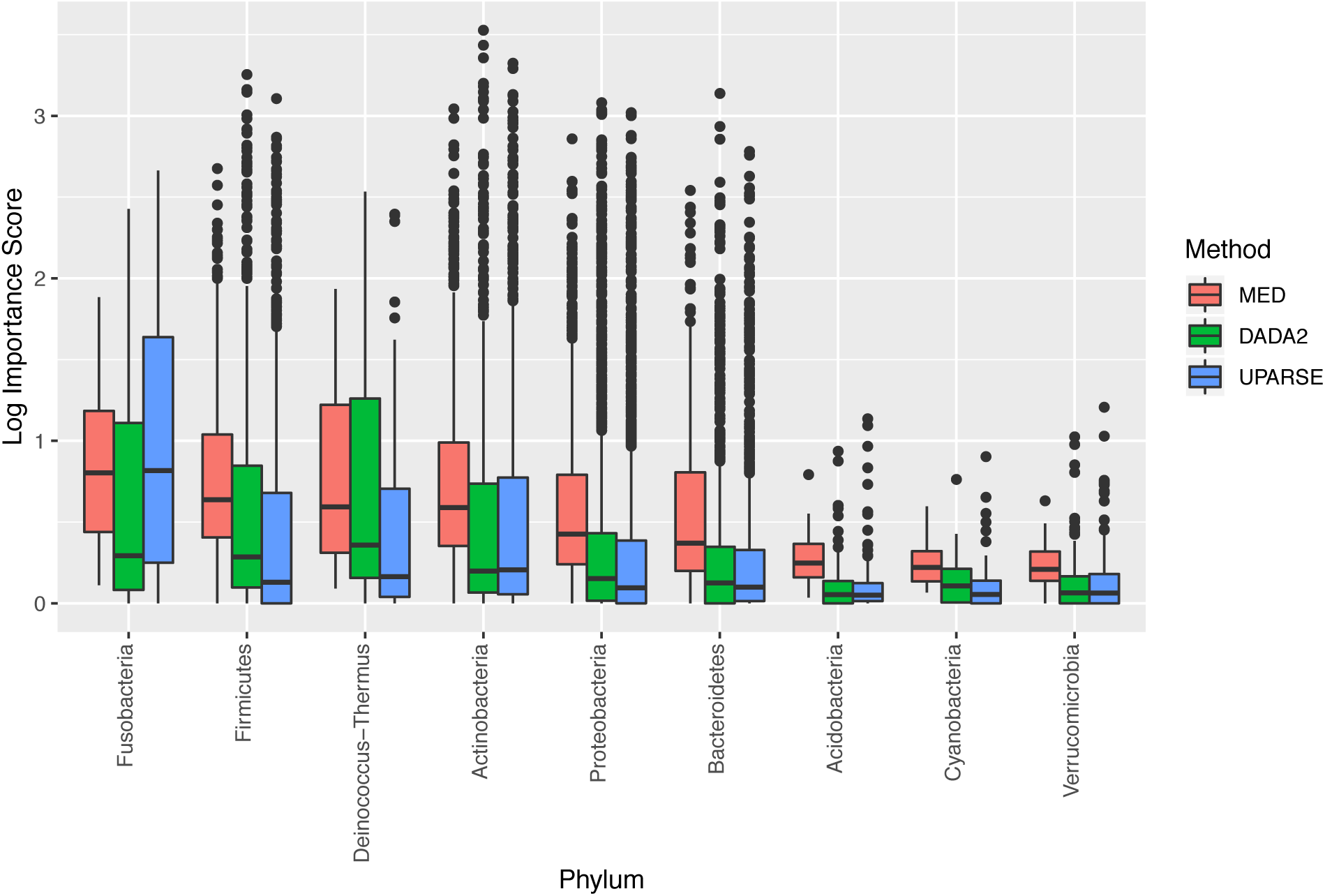
Distribution of importance scores by phylum and sequence processing method. Importance score is log transformed to aid visualization. All phyla are significantly different by Kruskal-Wallis test, and MED has a higher average importance score for all phyla except for Fusobacteria. There are a large number of outliers in abundant taxa, due to the non-normality of importance scores.

**Supplementary Figure 2:**
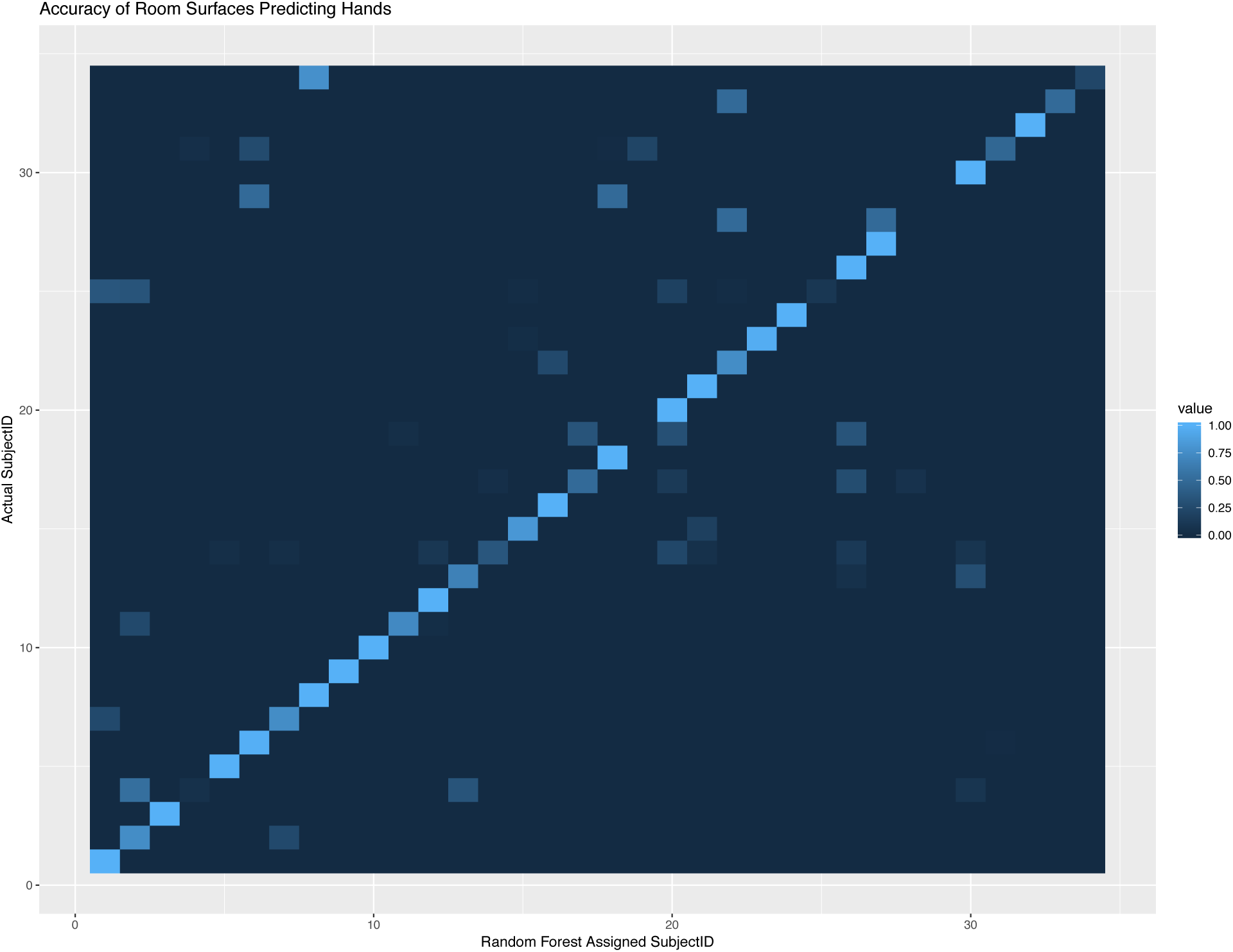
A confusion matrix generated by the results of a random forest. It compares the actual identity of a sample with the one assigned by the random forest. Accurate classification appears on the diagonal, and any deviation is a mislabeled sample. Mostly samples fall on the diagonal, reflecting the 4.76 error ratio.

**Supplementary Figure 3:**
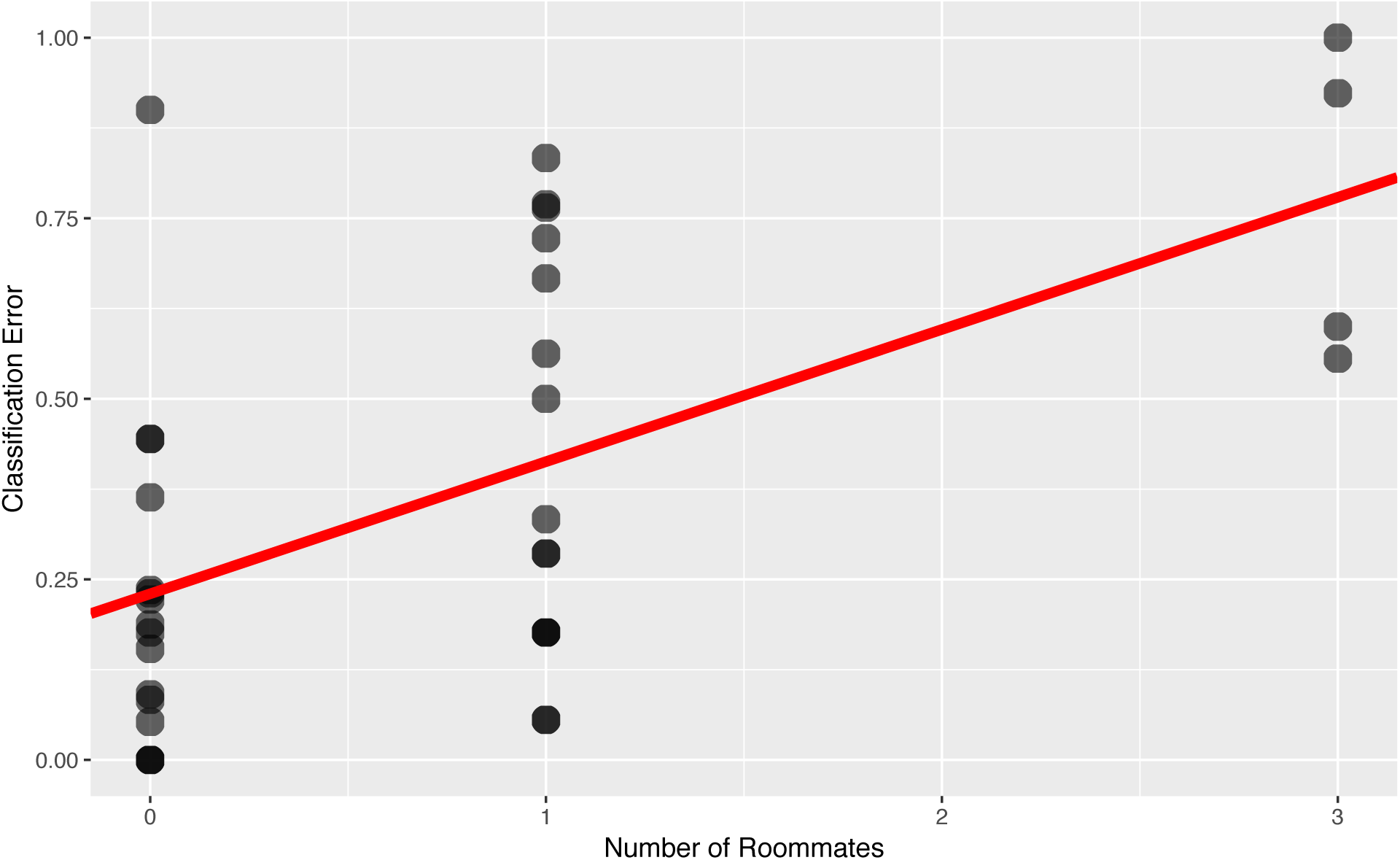
Classification error plotted against the number of roommates.

**Supplementary Figure 4:**
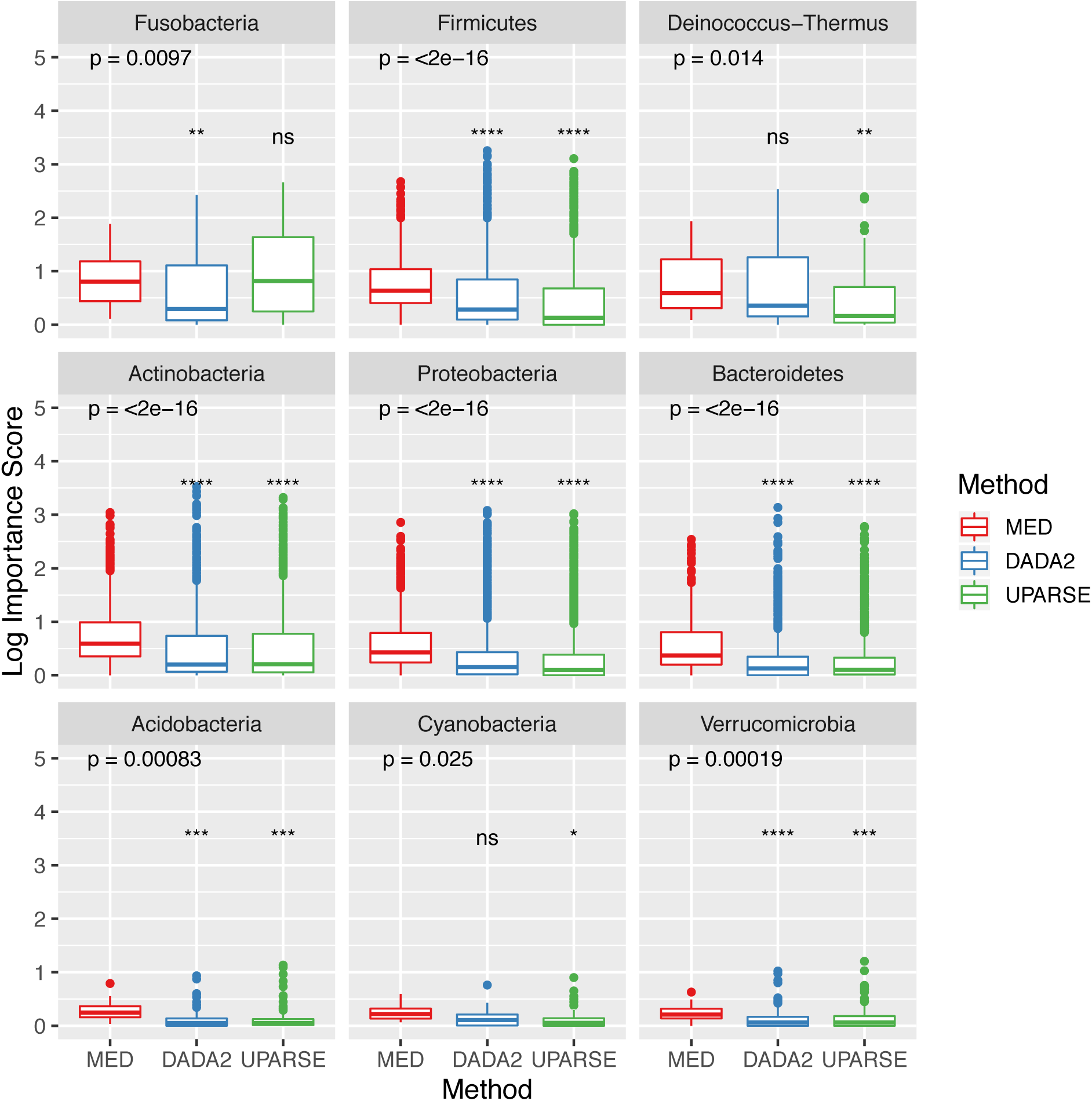
The distribution of importance scores by phylum, with both Kruskal-Wallis significance between all pairs, and Wilcox-results of means compared to MED importance scores

**Supplementary Figure 5:**
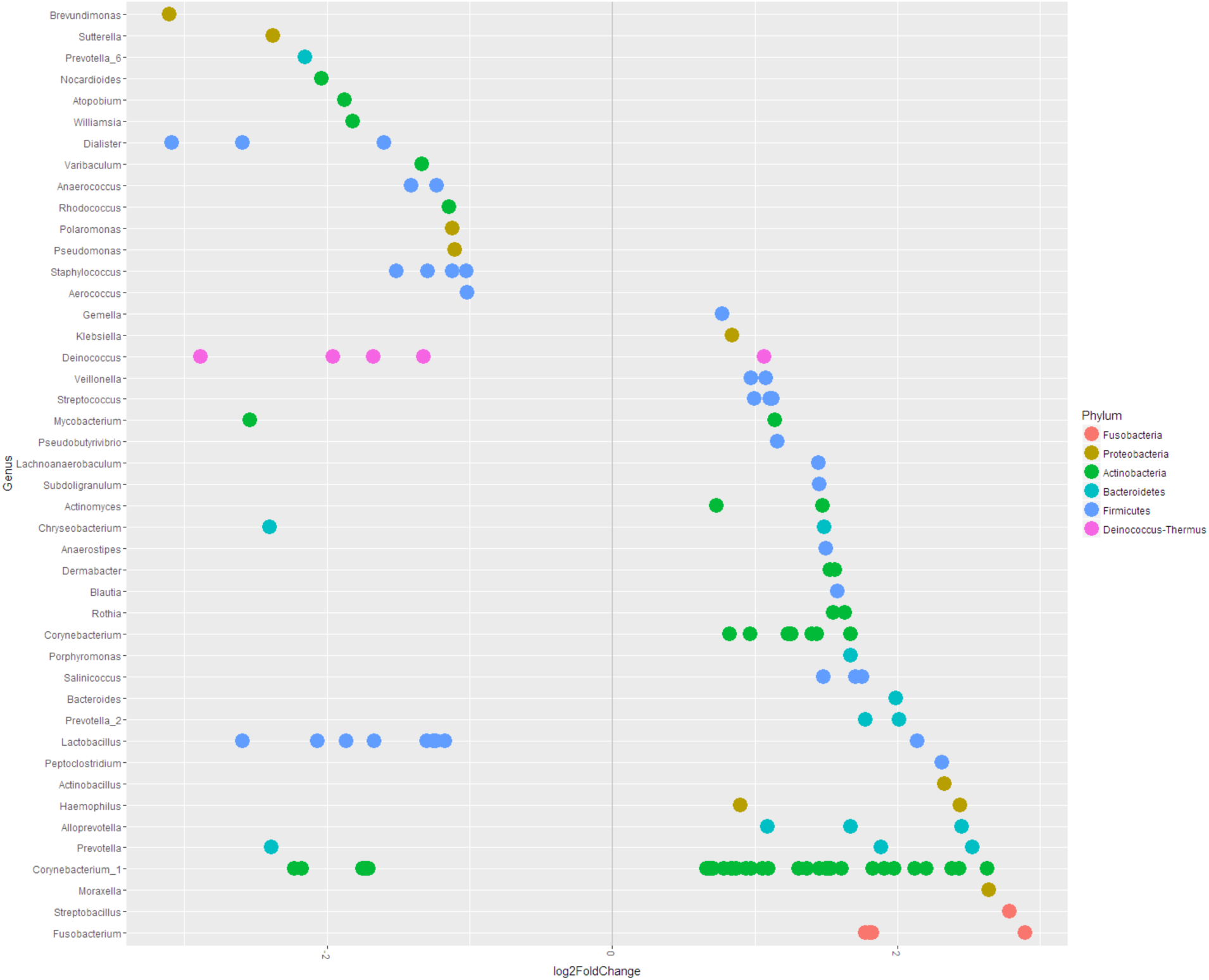
Differential abundant sequences between male and female individuals, with women at left and men at right.

**Supplementary Figure 6:**
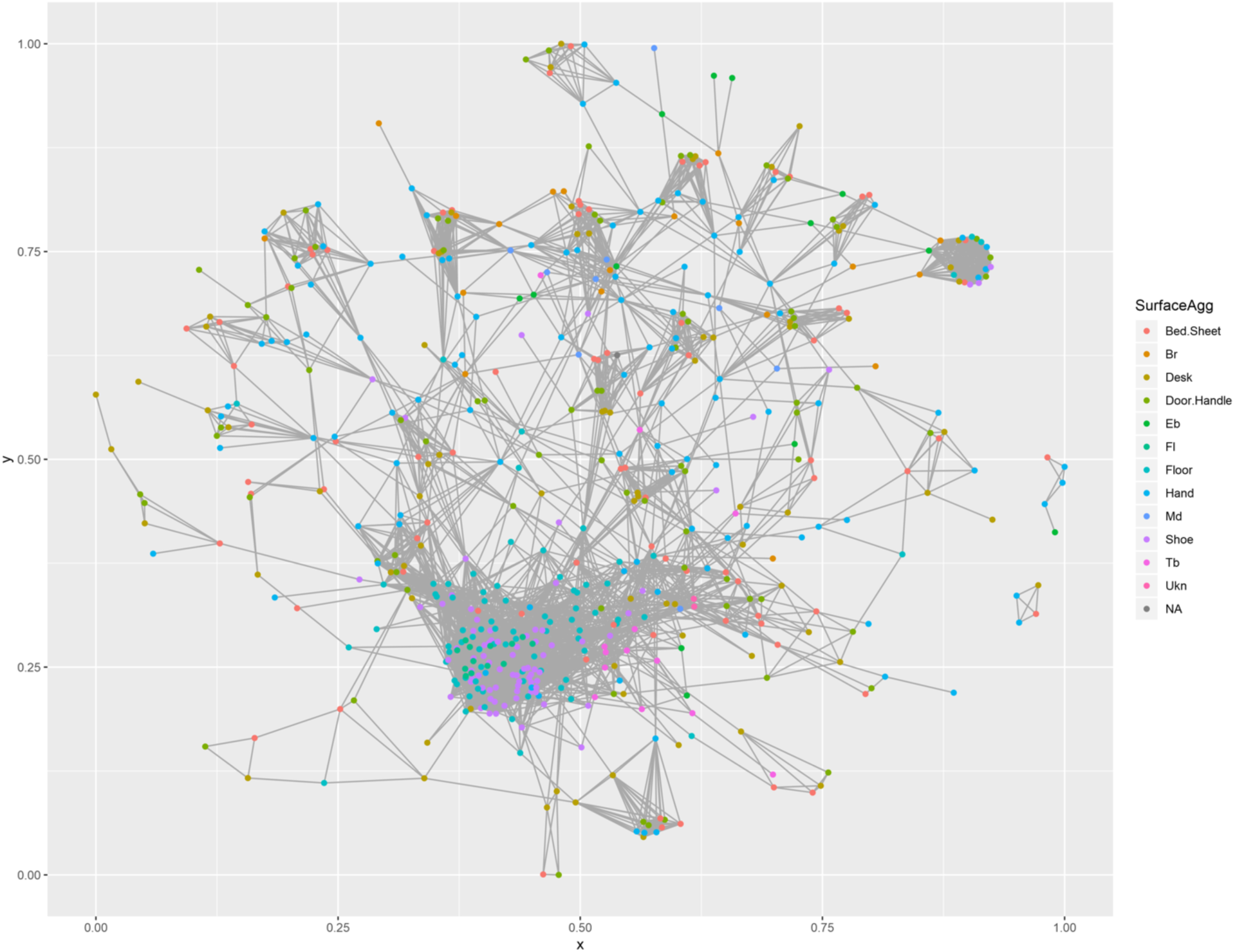
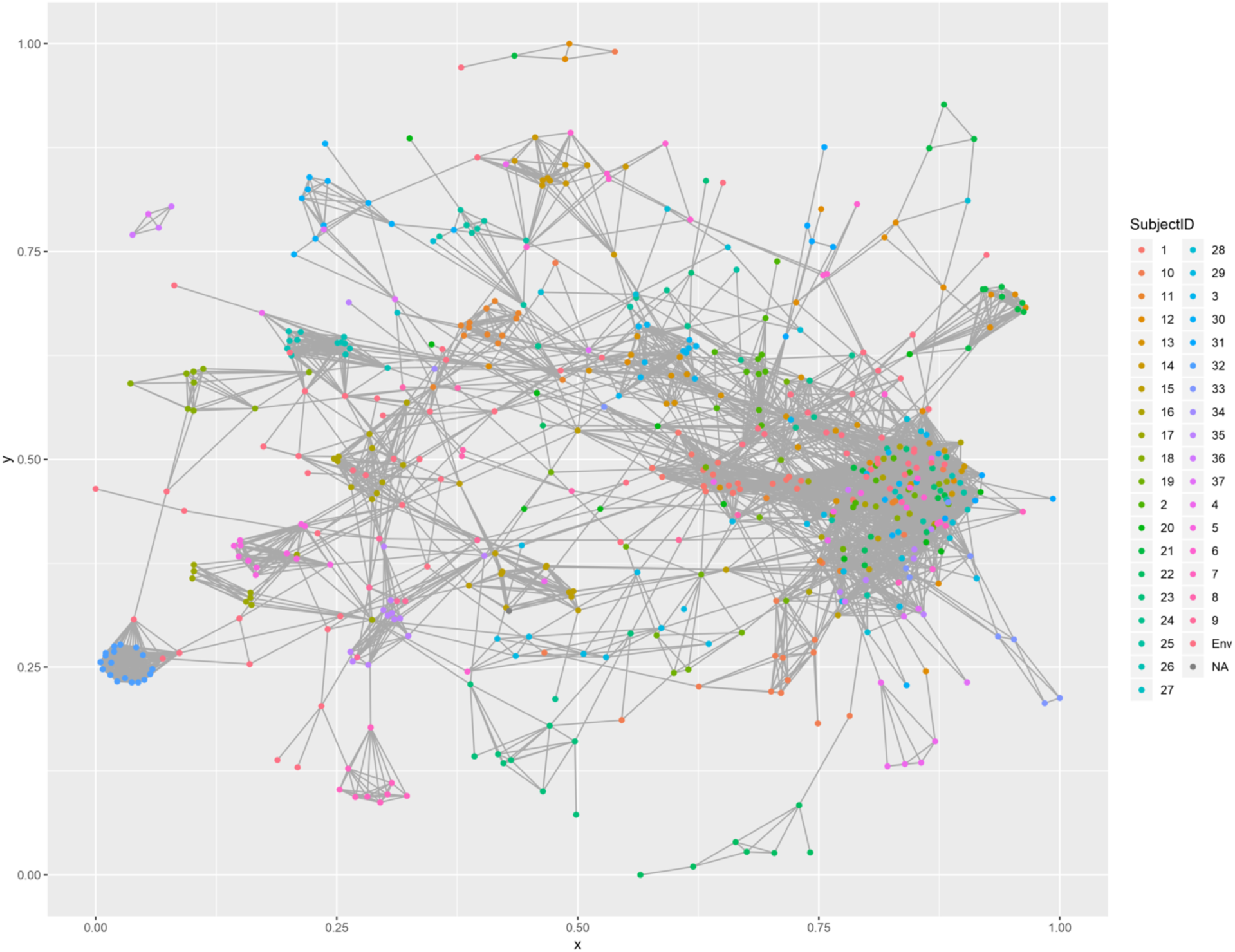
(a) Random forest proximity graph colored by individual, (b) colored by surface type.

**Supplementary Figure 7:**
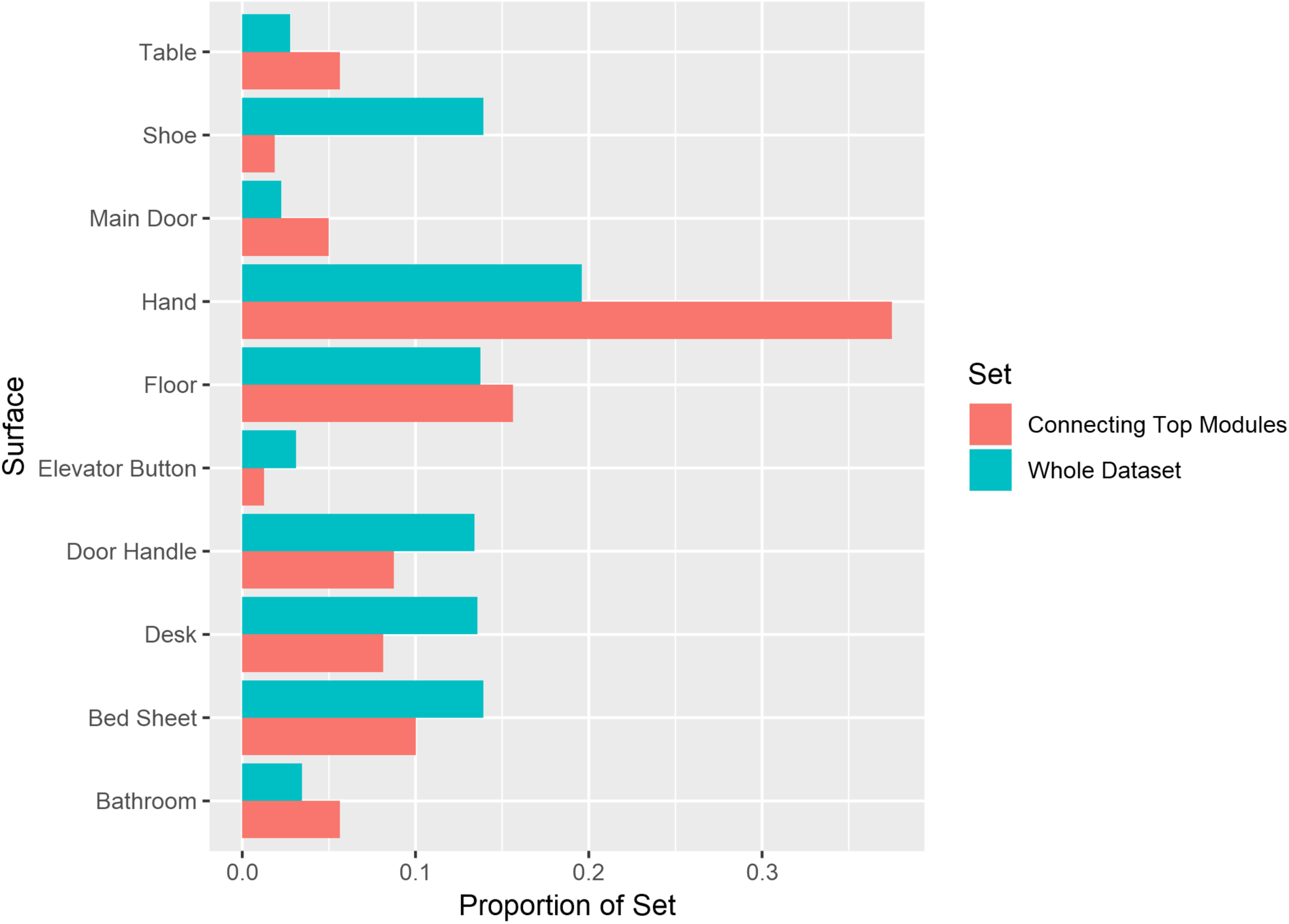
Bar graph comparing the proportions of samples connecting top modules compared to those in the dataset at large. Hands show significant enrichment in connecting modules, indicating that they are the likely source of exchange between modules.

